# Expressing glycan motifs as a containment order increases sensitivity and uncovers motif redistributions

**DOI:** 10.64898/2026.09.10.750563

**Authors:** Xinyu Zhao, Daniel Bojar

## Abstract

Glycan motifs are the standard readout of comparative glycomics, and pipelines treat them as independent features. Here, we show they are not. Motifs are partially ordered by substructure containment, so a parent motif’s abundance dominates each of its children. We make that order explicit as a directed acyclic graph, which stratifies the multiple-testing family correctly and, together with an empirical-Bayes variance prior taken from each motif’s containment neighborhood, raises estimated true positives across 45 glycomics datasets from 300 to 442 (+47%, p = 0.0001) after controlling for permutation-null false positives. Because a parent’s children and its residual form a genuine sub-composition, their logratio balances need no reference frame and no scale model, and they separate a motif’s own change from one inherited from its contexts. Of 337 significant parent motifs, 150 (45%) thus carry no signal of their own, while 138 motifs move only in the decomposition. We present case studies for both glycomics and glycoproteomics, including a recurrent reapportioning of core-1 sialylation at the immune-inhibitory disialyl-T antigen across eight human *O*-glycomes, and a colorectal fucosylation shift confined to the antenna, neither of which any marginal analysis reports.

## Introduction

Glycosylation is the most abundant and most structurally diverse co– and post-translational modification in biology, decorating over half of all proteins as well as lipids and the cell surface itself, and occurring as free glycans in mammalian milk [1–3]. Glycans mediate cell-cell recognition, immune regulation, and host-pathogen interaction [4–6], and aberrant glycosylation is a recognized hallmark of malignant transformation [7,8] as well as a rich source of clinical biomarkers [9].

What makes glycans analytically distinctive is that they are not templated. No gene encodes a glycan sequence, instead structures emerge from competition between glycosyltransferases and glycosidases for shared substrates in the secretory pathway [10,11], which produces a combinatorial repertoire far larger than any sample contains [12]. Biological function, however, is usually carried not by a whole structure but by a recurring substructure, a glycan motif. Examples include sialyl-Lewis X as the selectin ligand that initiates leukocyte rolling [13], the Tn and T antigens as tumor-associated epitopes with defined receptors [14], core fucosylation on IgG governing antibody-dependent cytotoxicity [15] and TGF-β receptor signaling [16], or bisecting GlcNAc modulating neural function [17]. Motif-level analysis is therefore not merely a dimensionality reduction but the level at which glycan biology is usually expressed, and it is the level at which comparative glycomics is routinely analyzed [18–20].

Since glycans are structurally degenerate, substructures such as terminal sialic acids, core-1 *O*-glycans, or Lewis epitopes, are quantified by summing the abundance of every structure that contains them, which can also be weighted by the number of their occurrence per glycan [19]. This turns an unwieldy and sparsely populated sequence space into a compact feature table of tens of interpretable terms, and that table is the universal intermediate on which differential abundance, classification, and biomarker discovery are all run.

Such a table is then typically analyzed as though its columns were independent measurements. They are not, and the dependence is not subtle. If a motif is contained in another, every structure counted toward the child is also counted toward the parent, so the parent’s abundance dominates the child’s in every sample and their test statistics are near-deterministically coupled. Across the 45 datasets analyzed here, a median of 61% of quantified motifs contain at least one other quantified motif, at a median graph depth of six. Independence is not a mild approximation in this setting, it is false for most of the table.

The Gene Ontology community met the same problem in the same shape and solved it explicitly. The elim and weight algorithms decorrelate the GO graph by removing the genes of significant child terms from their ancestors [21], the focus-level procedure combines closed testing with a graph-respecting selection to control the family-wise error rate [22], and hierarchical FDR performs the analogous procedure for false discovery [23]. Glycomics has no equivalent.

These methods do not transfer directly, for three reasons that also explain why the glycan case admits a stronger solution. A GO analysis inherits a fixed, expertly curated ontology, whereas the motif vocabulary is produced anew by every experiment from whatever structures were detected, so the graph must be derived from the data rather than looked up. GO enrichment asks whether a term is over-represented in a selected gene list, whereas motif analysis has continuous, compositionally constrained abundances for every term in every sample. And GO parent-child relations are set inclusions with no quantitative consequence, whereas motif containment forces an exact inequality on abundance in every sample. That last difference is what makes the decomposition below possible. A glycan parent and its children are numerically related, not merely conceptually nested, so the parent can be split into parts that sum to it.

Prior approaches to this general problem include GlyCompare [24], which orders substructures by biosynthetic precursor and reduces them to a minimal non-redundant set, and glycowork’s wildcard subsumption [19], which drops generalized motifs that carry no information beyond their specified counterparts. Both prune redundancy, yet neither builds a containment partial order, neither uses it for error control, and neither decomposes a parent’s change into its child contexts.

Here we do all three. We construct the containment order over whatever motif vocabulary an experiment produced, use it to borrow variance between motifs that contain one another, use it to stratify the testing family, and use it again to ask, for every motif with children, whether its change is its own or inherited. The last question turns out to be answerable without any reference frame or scale model at all, which in a compositional measurement [20] is unusual enough to be worth stating on its own. This leads to enhanced sensitivity, benchmarked against real-world glycomics data, and allows for new kinds of inquiries into motif dysregulation. With several real-world examples, we demonstrate how our new approach (fully contained in glycowork v1.10.0+) can lead to new biological insights from comparative glycomics data.

## Results

### The motif containment order is dense rather than sparse

Formally, the motifs of a glycomics dataset form a partially ordered set under containment (Fig. 1A). We write m ⊑ m′ when motif m is a substructure of motif m′. This relation is reflexive, antisymmetric, and transitive, but it is only partial. Many motif pairs are simply incomparable, so the motifs form neither a list nor a tree but a directed acyclic graph (DAG) in which a motif may have several parents and several children. What distinguishes containment from an abstract ordering is that it is enforced in the data. If m ⊑ m′, then every detected structure carrying m′ also carries m, so the abundance *a* of the two motifs obeys *a*_m_(*s*) ≥ *a*_m_′(*s*) in every sample *s*. The order is thus more than an annotation layered on top of the measurements, and rather represents a set of inequalities the measurements already satisfy.

**Figure 1.**
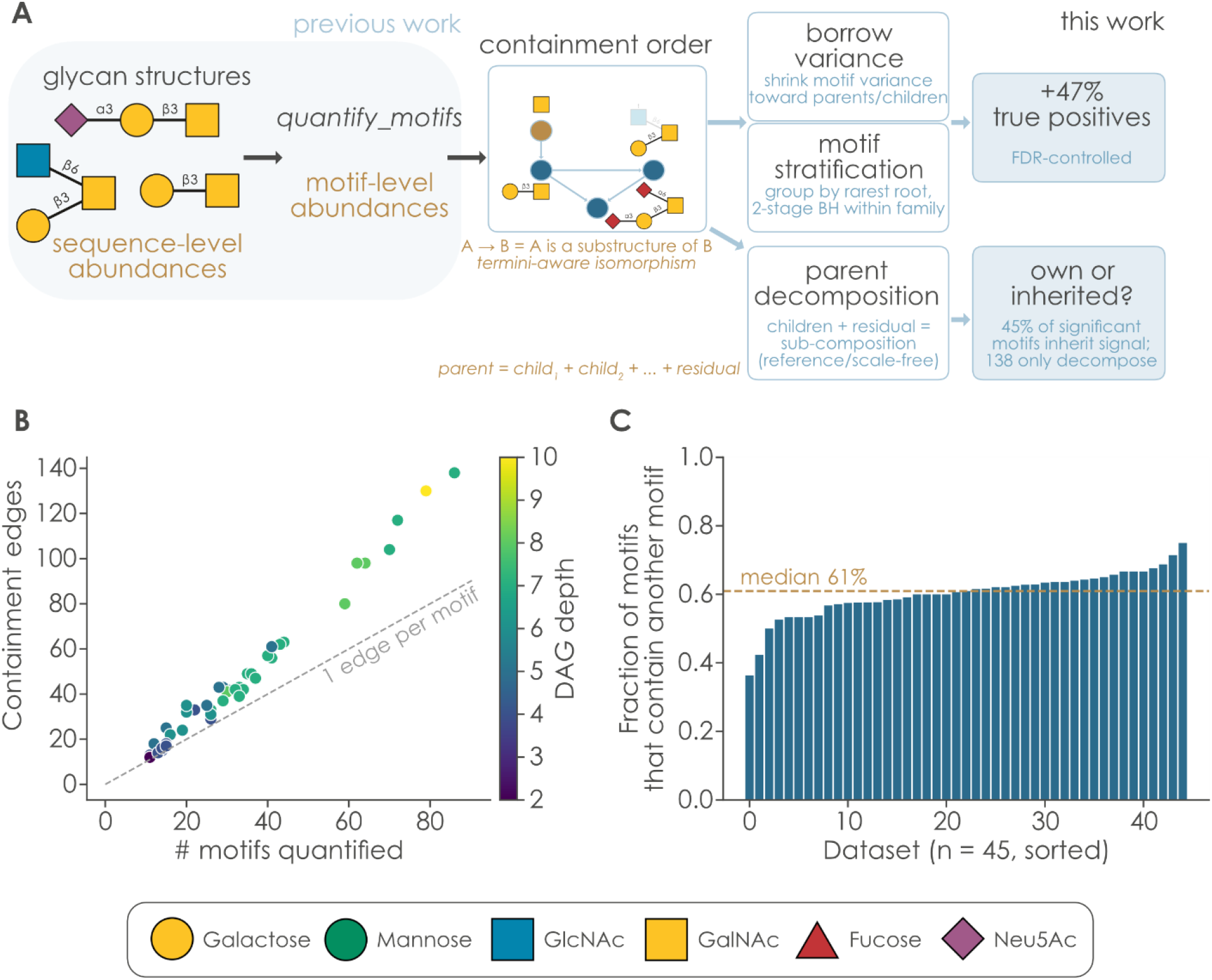
Glycan motifs are a containment order. **A)** Schematic of our workflow. Motifs are quantified via glycowork’s *quantify_motifs* function and ordered by containment. That single order is then put to three uses: to borrow variance across a motif’s parents and children when testing it, to stratify the multiple-testing family, and to decompose each parent into its child contexts. The first two raise detection sensitivity, the third separates a motif’s own change from an inherited one. In the graph inset, brown marks a root motif and blue its descendants; arrows point from a substructure to the structures that contain it. **B)** Every motif set carries more containment edges than it has motifs. Each point is one dataset (n = 45): quantified motifs on the x-axis against containment edges on the y-axis, filled by graph depth (viridis, dark = shallow). The dashed gray line marks one edge per motif; all datasets lie above it. **C)** A median of 61% of quantified motifs contain at least one other quantified motif. Per-dataset fraction of quantified motifs that contain at least one other quantified motif, sorted; the dashed brown line is the median (61%). All glycans in this work have been drawn using GlycoDraw [25] and are in line with the Symbol Nomenclature for Glycans (SNFG).

Two things follow. Statistically, a single global estimate of the proportion of true null results is thus spread across branches whose null proportions differ considerably, which unnecessarily costs statistical power. Biologically, a change in one structure is reported once for every ancestor that contains it, so a table showing significant decreases in sialic acid, in galactose, in core 1, and in sialyl-T antigen may be showing one event four times, and it gives the analyst no way to tell which of the four is the actual event.

We build this order directly from the motifs an experiment quantified, so that it adapts to the vocabulary rather than assuming a fixed hierarchy. An edge from motif A to motif B means A is a substructure of B, tested by termini-aware subgraph isomorphism within glycowork. A host structure may attach anything at a child’s open ends, so only what the child itself pins down is provable, and a node with something above it is internal while an open end is either the child’s declared terminus or genuinely unknown. Two design decisions make this tractable and unambiguous. Abundance dominance is an exact necessary condition for containment, since every structure counted toward B is also counted toward A, so it prefilters candidate pairs before any isomorphism is attempted. And when two labels are mutually isomorphic, the more ambiguous one, the one matching strictly more structures, becomes the ancestor, with a positional index as a total-order fallback that guarantees acyclicity. The graph is then transitively reduced so that each edge is an immediate containment.

Across the 45 glycomics dataset s curated within glycowork the resulting orders are dense (Fig. 1B). A median dataset quantifies 29 motifs joined by 39 containment edges, with 2,054 edges in total, a median depth of six and a median of four roots. The fraction of motifs that contain at least one other motif is 61% at the median and never falls below 38% (Fig. 1C). Grouping by rarest root ancestor partitions these into a median of four families with essentially nothing left over. Across all 45 datasets only seven motifs in total fall into the unassigned remainder.

The families are not arbitrary partitions but recognizable biosynthetic branches, discovered from the data rather than imposed (Supplementary Fig. 1). In a colorectal *O*-glycome the roots are Fuc, Neu5Ac, GlcNAc, GalNAc, GlcNAc6S, and GalOS, and each collects exactly the epitopes built on that residue. The Fuc family holds the A, B, and H antigens and the Lewis structures, the Neu5Ac family holds sialyl-Lewis A, disialyl-T, CAD, and Sd^a^, and the GlcNAc6S family holds the keratan-sulfate-type structures. In an *N*-glycome the roots are instead, e.g., HexNAc, Sia, Hex, Fuc, and GalOS, with the Hex family carrying the high-mannose and *N*-glycan core mannose motifs and the Fuc family separating core fucose from the antennary Lewis A/X epitopes. Because each family is a pathway branch, giving it its own null proportion is not merely a statistical device but the assumption that different arms of glycan biosynthesis respond independently.

### The containment order recovers signal that a flat correction discards

We assigned each motif to the family of its rarest root ancestor and applied two-stage adaptive Benjamini–Hochberg multiple testing correction within families, so that each branch received its own estimate of the proportion of true nulls. We then benchmarked this against the same pipeline with a flat correction (i.e., glycowork v1.9.0). Both arms shared identical preprocessing, identical test statistics, and identical effect-size gates, with the only difference being the leveraging of motif containment for test stratification and variance borrowing. Because a more permissive correction can always manufacture more hits, we measured sensitivity as the number of significant hits above an effect-size gate minus the same count under 100 label permutations of the same data by the same method. The permutation null mean then represents that method’s own false-positive count, so the subtraction can penalize each method for its own liberality before the two are compared.

Because of the variance borrowing, the order also improves the test statistic itself, not only the correction. Glycomics cohorts are small, with a median of 14 samples here, so a per-motif variance estimated from a handful of replicates is unstable, and an underestimate produces a spuriously large t or F. The standard remedy is empirical-Bayes moderation, in which each feature’s variance is shrunk toward a prior learned across features [26]. Containment supplies a better prior than the usual global one, as motifs that contain one another are measured on overlapping sets of structures and therefore share measurement noise, so a motif’s parents and children are a more relevant variance reference than the rest of the table. We therefore shrink each motif’s within-group variance toward the geometric mean over its own containment neighborhood, with the prior degrees of freedom estimated per dataset by Smyth’s moment estimator and capped at the residual degrees of freedom so that the prior can never outweigh the data, bounding this hyperparameter following the rationale of Phipson et al. [27]. Across the corpus, this neighborhood prior outperformed a global prior in the two-group designs at every shrinkage setting we examined, and because the moderation enters only the p-value, all reported effect sizes remain the ordinary unmoderated Cohen’s *d* and ω^2^.

Combining the stratified correction with the neighborhood variance prior recovered substantially more signal across the corpus (Fig. 2A). Estimated true positives rose from 299.9 to 441.6 across all 45 datasets, a 47% gain that was higher in 30 of 45 datasets (stratified sign-flip test across designs, p = 0.0001) and that survived deletion of any single dataset (p < 0.05 in all 45 leave-one-out replicates, worst case p = 0.0003). Both designs gained significantly on their own, +67.2 in the 30 two-group datasets (p = 0.0078) and +74.5 in the 15 multi-group datasets (p = 0.0044). Sweeping the lower effect size gate from zero to the top of its range left the advantage positive throughout (Fig. 2B), and repeating the whole benchmark with ceilings from 2 to 50 gave pooled gains between +118 and +142 at p < 0.007 in every case, so neither bound of the window drives our result. This increase in sensitivity resulted in roughly ten true positives for each additional false one, and the estimated false-positive share of all hits was essentially unchanged (∼5%; Fig. 2C).

**Figure 2.**
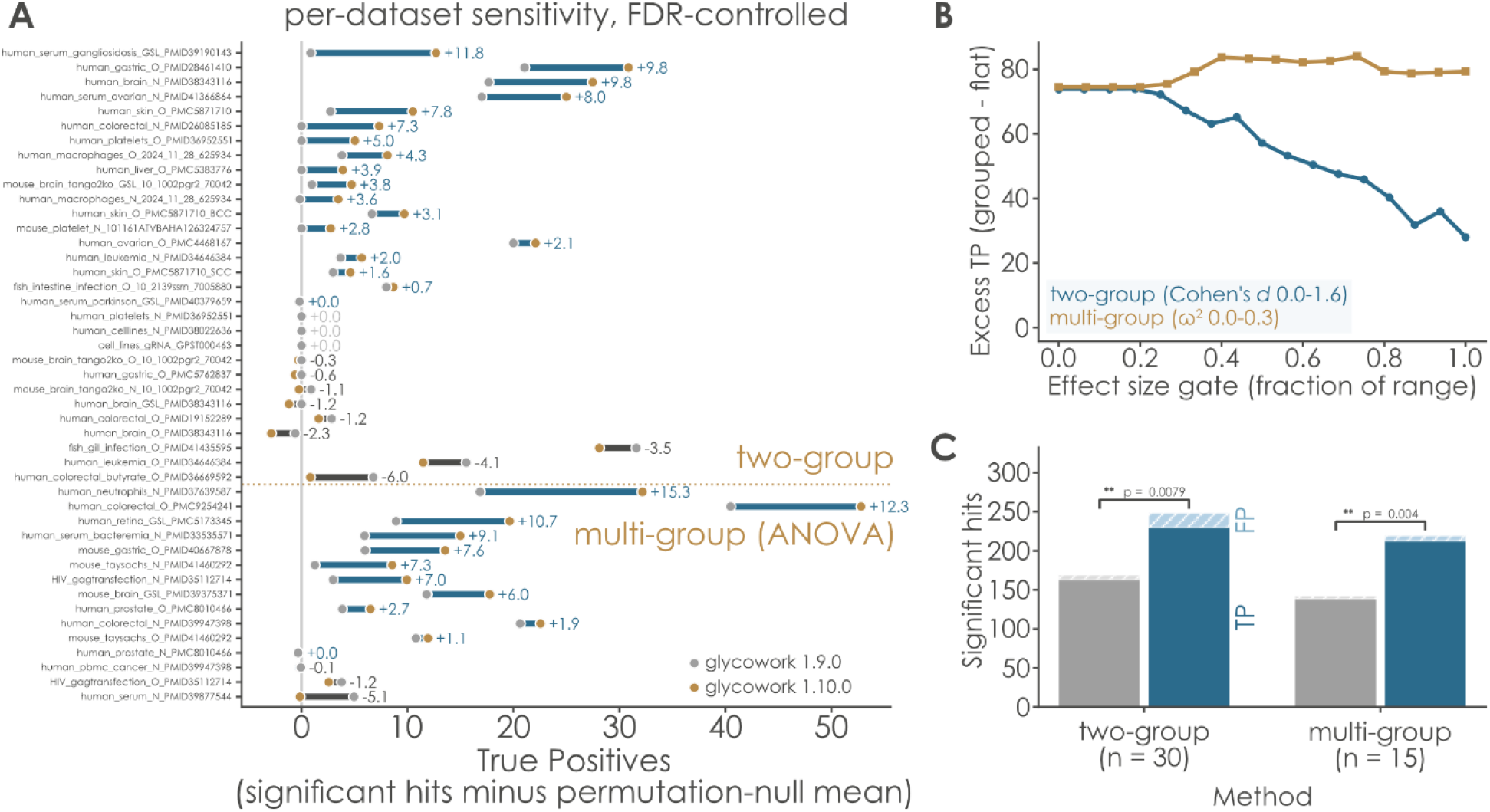
Treating motifs as a containment order raises detection sensitivity. **A)** The containment-aware pipeline estimates more true positives in 30 of 45 datasets. Each row is one dataset, split into two-group and multi-group blocks by the dotted line. Gray circles, flat Benjamini-Hochberg and no variance moderation (glycowork v1.9.0); brown circles, DAG-grouped correction and empirical-Bayes variance moderation (v1.10.0). The connecting segment is blue where the grouped correction estimates more true positives and black where it estimates fewer, and the number at the end of each segment is the difference. The x-axis is significant hits above the effect-size gate minus that same method’s mean count over 100 label permutations. **B)** Both designs gain at every effect-size gate. Summed difference in estimated true positives (grouped minus flat) as the lower effect-size gate is swept across its range with the ceiling of |*d*| ≤ 5 held throughout, blue circles for two-group and brown squares for multi-group tests. **C)** Containment-aware analysis returns roughly ten additional true positives for each additional false one. All significant hits pooled per test type, with the solid portion of each bar the estimated true positives and the hatched portion the estimated false positives (the permutation-null mean); gray, flat correction; blue, DAG-grouped. Brackets give the sign-flip permutation test on per-dataset estimated true positives. Throughout, hits are counted within the window 0.5 ≤ |*d*| ≤ 5 for two-group and ω^2^ ≥ 0.06 for multi-group comparisons.

A paired six-versus-six gastric *O*-glycome illustrates why the upper bound is needed. Without it, v1.9.0 returns 24 hits there with Cohen’s *d* values between 14 and 29, and its own permutation null reproduces up to 24 such hits under randomized labels, so the apparent signal is a variance collapse that survives label scrambling. v1.10.0 calls nothing in that dataset significant, correctly, since its smallest corrected p-value is 0.24 across 64 hypotheses at six samples per group. Uncapped, this single dataset contributes an apparent loss of 23.9 estimated true positives to the older pipeline’s credit; inside the window it contributes 0.6, and the three other datasets in the negative tail behave the same way.

Three datasets moved from no recoverable signal at all to 26.9, 10.9, and 3.5 estimated true positives. In the human liver *O*-glycome (n = 6) the new pipeline surfaced a coordinated gain of Lewis X, with internal Lewis X up 2.2 log_2_FC units and terminal Lewis X up 2.7; the decomposition (detailed below) further showed this to be a genuine conversion rather than a bulk increase, as internal LacNAc redistributed (p = 0.007) while terminal LacNAc lost share (p = 0.019). In human cell-line *N*-glycomics (n = 6) it surfaced a wholesale gain of Lewis-type epitopes in THP-1 cells versus MDA-MB-231, with terminal Lewis A up 15.5 log_2_FC units and sialyl-Lewis X up 13.0, alongside a redistribution of GlcNAcβ1-4Man (p = 4.2e-07). Both are conclusions that a flat correction returned as an empty table.

We compare our new approach here to the flat correction the current state-of-the-art method for comparative glycomics analysis uses, which is the relevant baseline for the claim that ordering the motifs increases statistical power, and it should not be read as a contest against the GO-derived procedures which address a different problem. GlyCompare here is not a competing correction, as it orders substructures by biosynthetic precursor and returns a minimal non-redundant set, so it removes redundancy before testing, where we retain it and model it. Our decomposition approach below is closer in spirit to topGO’s elim, which tests a parent after removing what its significant children explain, except that here the subtraction is exact and quantitative rather than a heuristic over gene lists, which could not be applied to glycomics data.

### Nearly half of all significant motif findings are inherited rather than their own

For a motif with children, we define the residual as the parent’s abundance minus the summed abundance of its children, i.e., the part of the parent that occurs outside every child context.

Children and residual sum to the parent, so they are a genuine sub-composition of it, and subtracting the log of any one part from the others returns that sub-composition’s additive logratios. Any per-sample scalar (e.g., total ion current, dilution, normalization constant, unmeasured biomass) cancels from a parent-to-child logratio. These balances therefore carry no reference frame and no scale model, which no other output of a compositional glycomics pipeline can claim. We then used a Hotelling’s T^2^ test on the balances (two groups) or a one-way PERMANOVA (multiple groups) to ask whether the parent redistributed across its child contexts. The residual is tested separately here, in the same logratio frame the marginal statistics were computed in, and answers whether the parent changed outside all of them. Redistribution, residual, and marginal tests answer different questions, so each is corrected as its own family. A parent that occurs only inside its children has no context of its own left to test and is recorded as such. Overall, coverage is high as 795 of 847 parent motifs (94%) received a redistribution test in this manner, 91% in two-group designs and 99.7% in multi-group designs. We note that all these analyses could be run automatically in glycowork v1.10.0+ in seconds (for typical datasets) and are present in the standard output of *get_differential_expression* and *get_glycanova* (if motif-analysis is chosen), for users to inspect.

Across the 45 datasets, 847 motifs had children and were therefore decomposable, and 337 of those were significant under the standard marginal analysis. Of those 337, 150 (45%) had a residual p-value of exactly one with a residual effect size of exactly zero, as the motif occurred only inside its children, and the entire marginal result was inherited from them (Fig. 3A-B). Another 145 had a genuine residual change of their own and 42 were marginally significant without the decomposition resolving which. The motifs most often inherited were precisely the ones the literature reports most often (Gal in 17 datasets, Galβ1-4GlcNAc in 14, Neu5Ac in 12, Galβ1-3GalNAc in 10, GalNAc in 8). These represent monosaccharide-level statements of the form “sialylation is increased” or “galactosylation is decreased” that are the common currency of the field, yet nearly always would have a more specific phrasing (and insight) that is missed in the reporting of these inherited effects. The fraction of inherited calls varied from a quarter to all of them across datasets, and was not confined to any one glycan class or tissue.

**Figure 3.**
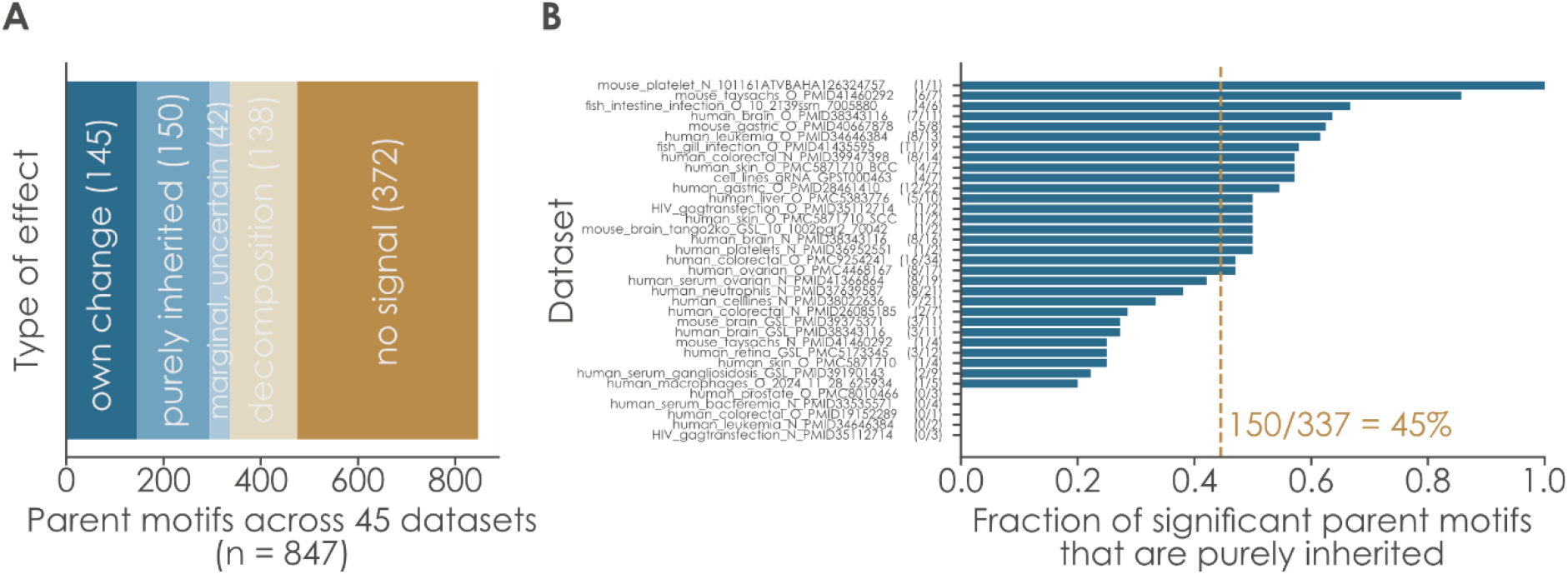
Most motif changes are redistributions, not marginal changes. **A)** Of 337 marginally significant parent motifs, 150 carry no signal of their own, while 138 further motif changes are visible only in the decomposition. All 847 parent motifs pooled across 45 datasets, classified by what the decomposition says: significant residual change of the motif’s own, marginally significant but purely inherited (residual p = 1 and residual effect size 0), marginally significant but unresolved, significant only in the decomposition and invisible marginally, or no signal. **B)** Every dataset carries inherited calls, ranging from a quarter to all of its significant parents. For each dataset, the fraction of its marginally significant parent motifs that are purely inherited; counts are given after each dataset name and the dashed line marks the pooled value (150/337 = 45%).

Two structural regularities became apparent in our analyses. First, inheritance is a property of branch points and not of nesting as such. Among significant parents with a single child, none of 120 is purely inherited, whereas 66% of those with two children are, rising to 79% at four or more (Spearman ρ = 0.62, p = 5.9e-37). A parent whose entire abundance sits inside one child is by definition the same feature and is removed at deduplication during preprocessing, so what the residual detects here is specifically the loss of a motif’s own context when its occurrences fan out across several alternatives. Second, the probability that a motif’s change is its own tracks how specific the motif is. Among motifs significant in at least four datasets, those never purely inherited are the linkage– and architecture-defined terms (e.g., Oglycan_core1, Oglycan_core2, Nglycan_complex, GlcNAcβ1-2/4/6Man, Neu5Acα2-3Gal, Neu5Acα2-6GalNAc, internal and terminal LacNAc; common name terms are from glycowork’s “known” feature set that aggregates literature-known motifs). Conversely, those always purely inherited are the monosaccharide-level terms Gal, GlcNAc, GalNAc, Galβ1-3GalNAc, and Galβ1-4GlcNAc. Therefore, the more precisely a motif is specified, the more likely that a change attributed to it is really its own.

### The decomposition sees 138 changes the marginal test cannot

On the other side of our analyses, 138 parent motifs across 25 datasets were not significant marginally, yet were significant in redistribution or residual. In other words, the motif’s total abundance did not move, but the way it was distributed over its sequence contexts did. These concentrated on sialylation, fucosylation, and the *N*-glycan core (Neu5Ac in 8 datasets, Fuc in 7, Neu5Acα2-3Gal in 7, Manα1-3Man in 6, internal LacNAc in 6), nodes that are close to universal and therefore almost never marginally interesting.

Redistribution also recurred across tissue and disease (Fig. 4). Among motifs testable in at least eight datasets, Neu5Ac redistributed in 20 of 43, Neu5Acα2-3Gal in 19 of 32, internal LacNAc in 16 of 30, Gal in 15 of 23, Fuc in 14 of 21, and Galβ1-4GlcNAc in 11 of 31. Among the 16 most recurrent parents there were 174 significant redistribution events, and 66 of them (38%) occurred in a dataset where that motif was not marginally significant at all.

**Figure 4.**
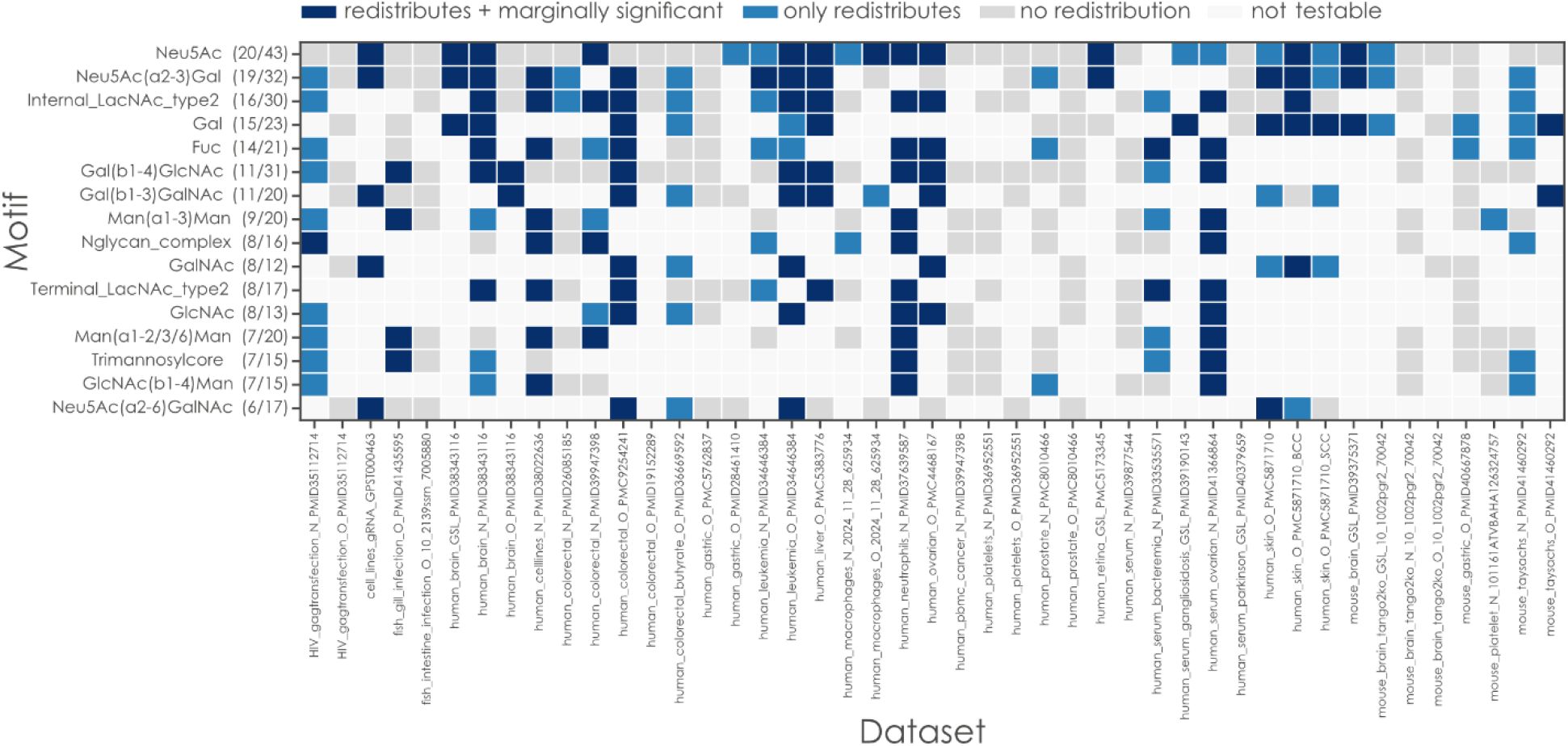
Decomposition tests identify motif redistributions. Rows are the 16 most recurrent parent motifs, annotated with the number of datasets in which they redistribute out of the number in which they are testable; columns are the 45 datasets. Dark blue, the parent redistributes significantly and is also marginally significant; light blue, it redistributes but is marginally invisible; light gray, testable with no significant redistribution; white, not testable in that dataset (the motif is absent or has no children there).

Redistribution detection improved only weakly with sample size and clearly did not require it, as 38% of parents redistributed significantly in datasets with eight or fewer samples, against 44% in those above twenty (Spearman ρ = 0.11, p = 2.8e-03). Because the balances we investigate here are ratios within a parent, much of the between-sample variation that limits marginal testing cancels out, which is why the decomposition remains informative in the small cohorts typical of glycomics.

Where redistribution occurs is itself informative (Supplementary Fig. 2). Glycosphingolipid motif sets redistribute in 44 of 67 testable parents (66%), well above *N*-glycans (160 of 341, 47%) or *O*-glycans (136 of 387, 35%; Fisher p = 9.2e-05 for GSL against the rest). Ganglioside biosynthesis runs as parallel a-, b-, and c-series, in which the same promiscuous enzymes elongate every series in lockstep, which predicts strictly linear precursor-product chains rather than the branch-point combinatorics of *N*-glycan antennae, and therefore predicts exactly this excess of nesting. With six GSL datasets, we offer this as a hypothesis rather than an established contrast. Redistribution is also not a disease phenomenon at all, as malignant and non-malignant datasets redistribute at an identical 43% of parents (Fisher p = 1.0), so reorganization of motif context is a general property of glycomes rather than something disease creates.

The most recurrent single node in the corpus was the disialyl-T antigen. Across eight human *O*-glycome datasets it was the child that captured redistribution at three different parents, core 1, Neu5Acα2-3Gal, and Neu5Acα2-6GalNAc, in 19 significant events (Supplementary Figure 2c). Its direction, however, was not consistent: it gained share in 8 of those events and lost share in the remainder, draining in skin, liver, and colorectal tissue, while accumulating in leukemia and prostate. Disialyl core 1 is built from sialyl-T antigen by the redundant enzyme pair ST6GalNAc-III and –IV, which must both be deleted before the antigen is lost [28], and it is the specific ligand for the inhibitory receptor Siglec-7 [29,30], so the balance between disialyl-T and its monosialyl precursor tunes an immune-inhibitory terminal epitope in either direction. The draining direction mirrors the classic observation that malignant colonic epithelium loses disialyl structures in favor of their monosialyl precursors through loss of an α2-6 capping sialyltransferase [31], although the documented event is on the type-1 chain rather than on core 1, so ours is an independent recapitulation of the same logic at a different branch of the containment order. Where disialyl-T drained in colorectal tissue, sialyl-Lewis A and the CAD/Sd^a^ epitope gained share instead. We note that this CAD/Sd^a^ gain is explicitly a share within a sialylated parent and is not evidence of an absolute increase, which would be in contrast to literature reporting Sd^a^ loss in colorectal cancer [32–34].

Next, we extended our redistribution analysis to site-level glycoproteomics, where we replace substructure containment by component-wise containment of glycan compositions. In other words, within the same glycosite, a glycoform contains another only if it dominates it in every monosaccharide, which again forces the exact abundance inequality our method needs (Supplementary Fig. 3A). On 326 glycoforms across 55 sites of the human milk *N*-glycoproteome, this yielded 326 composition nodes joined by 168 containment edges at depth four, with 86 decomposable parents. Because compositions are ordered only within a glycosite, glycoforms on different proteins never contain one another, and the resulting order is shallower and sparser than any released glycome in this study (0.51 edges per node against a median of 1.37). Even so, the decomposition remains informative: comparing colostrum with mature milk, 36 of 196 tested glycoforms were significant in redistribution or residual, among them the IGHA1 and TRFL glycoforms whose composition shifted while their glycosite totals did not. Sparser containment therefore does not disable the decomposition, it merely narrows how much of the glycoform table is decomposable at all.

In another glycoproteomic dataset, of blood from healthy and gastric cancer individuals, a total of 11,638 site-specific *N*-glycoforms were identified across 1,295 *N*-glycosites [35]. Among them, 238 decomposable parent glycoforms from 67 *N*-glycosites were identified using our algorithm (Supplementary Fig. 3B). In particular, the glycoform “ZA2G-Asn112-H5N4A2” showed no significance in the marginal analysis but exhibited redistribution among its children, H6N5A2 and H6N5A3F1.

### Case studies

In basal cell carcinoma, flat motif analysis of the skin *O*-glycome returned 12 significant motifs, among them Neu5Ac, Gal, Galβ1-3GalNAc, and GalNAc, the monosaccharide-level terms a study would ordinarily report (Fig. 5A). All four had a residual p-value of exactly one. None of them changed on its own, each moved because the structures containing it did. What did change on its own was narrower and more specific: Neu5Acα2-3Gal (residual p = 1.9e-04, *d* = +1.17), core 1 (residual p = 2.4e-04, *d* = +1.08), internal LacNAc (residual p = 9.7e-04, *d* = +0.87), and α2-6-sialylated GalNAc, which fell (residual p = 8.5e-04, *d* = –0.91) despite not being marginally significant at all. Möginger et al. (2018) [36], analyzing the same cohort structure by structure, reported that total sialic acid, fucosylation, and sulfation were unchanged while α2-3-Neu5Ac increased and α2-6-Neu5Ac decreased, the latter found exclusively on core 1. Our decomposition recovers precisely that opposition of the two sialyl linkages automatically from the motif table, including the α2-6 loss that the marginal test misses entirely, whereas a flat analysis of the same table reports only the inherited monosaccharide terms.

**Figure 5.**
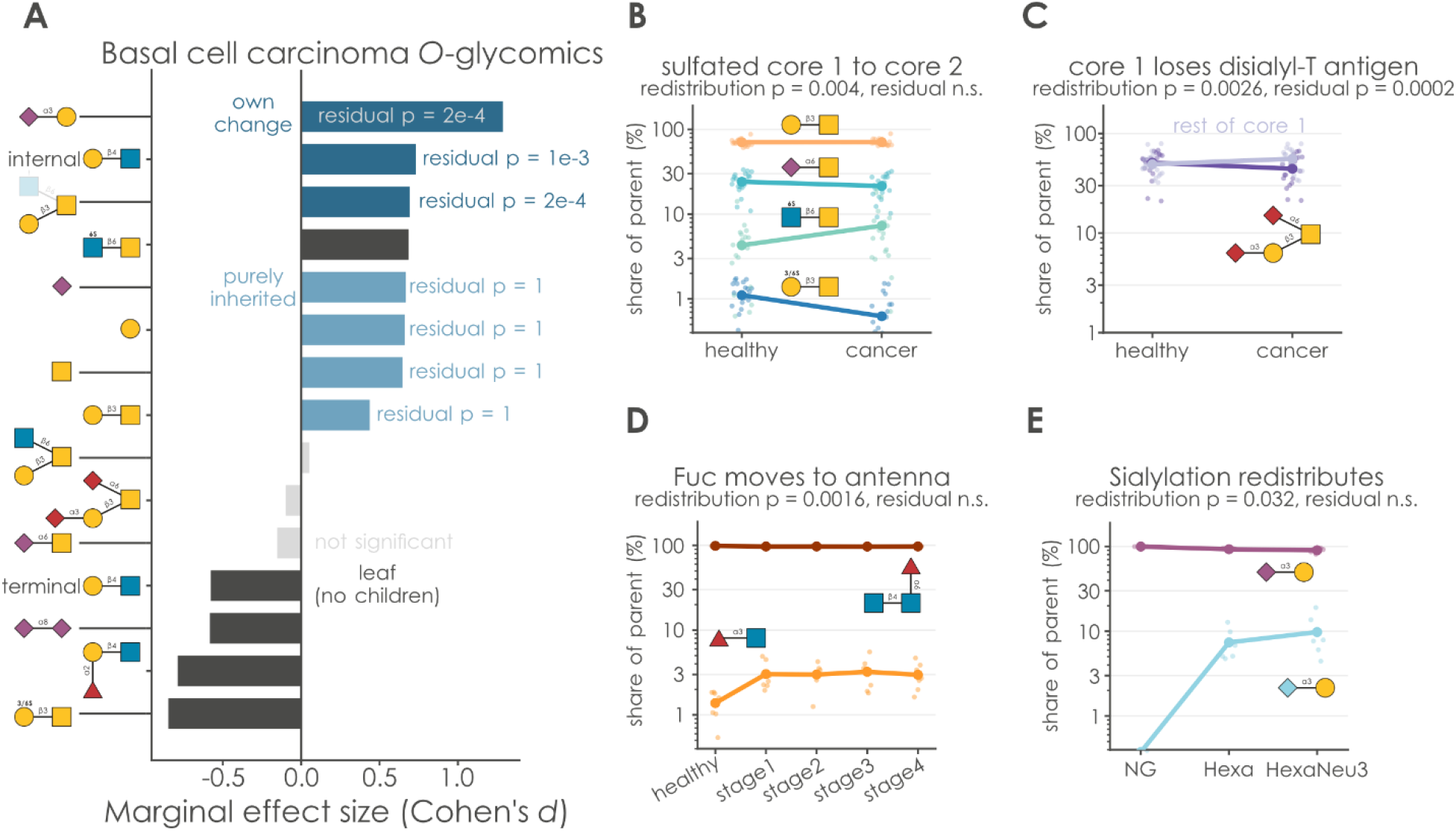
Uncovering new biology with motifs as a containment order. **A)** In basal cell carcinoma, the monosaccharide-level motifs carry no signal of their own. Basal cell carcinoma versus healthy skin: every quantified motif, ordered by marginal effect size and colored by decomposition verdict, light blue, purely inherited; dark blue, a significant residual change of its own; dark grey, a leaf with no children to decompose; light grey, not marginally significant. Residual p-values are printed beside the colored bars. **B-E)** Parent motifs reapportion across their child contexts while their totals stay flat. Child shares of a parent motif across groups. Each colored series is one child or the residual, small translucent points are individual samples and the connected large points are group means; the y-axis is the percentage of the parent carried by that part, on a log scale because shares span two orders of magnitude. Legends give the mean share in the first and last group. Panels: (B) GalNAc and (C) core 1 in basal cell carcinoma, (D) fucose across five colorectal cancer stages, (E) α2-3 sialylation across *Hexa* genotypes. Redistribution and residual p-values are given above each panel; * p < 0.05, ** p < 0.01, *** p < 0.001.

The redistribution itself is legible in the underlying shares within this dataset (Fig. 5B-C). Within GalNAc, the sulfated GlcNAc6Sβ1-6GalNAc context grew from 4% to 7% of the parent while α2-6-sialylated GalNAc fell from 24% to 21%, with the parent total essentially flat (redistribution p = 0.004, residual p = 1). Within core 1, the disialyl-T antigen share fell from 51% to 44% (redistribution p = 0.003), and here the residual was significant too, so core 1 both shrank and rearranged, which is an important conclusion that is difficult to reach with existing methods.

In colorectal cancer *N*-glycomics across five stages, total fucosylation never became significant, yet Fuc redistributed strongly (Fig. 5D). Antennary Fucα1-3GlcNAc rose from 1% to 3% of all fucose between healthy tissue and stage 1 and remained there, while core fucose was flat. A tripling of the Lewis-type branch that left the total unmoved was invisible to any marginal test of total fucosylation, and the independence of core (FUT8-mediated, α1-6) from antennary and Lewis-type (FUT3/4/6/7, α1-3) fucosylation is documented in colorectal cancer, where linkage-resolved profiling of 25 cell lines separated core from antennary fucosylation and tied them to different phenotypes [37], while FUT8 knockdown remodels antennary fucosylation without abolishing it [38].

Finally, in Hexa-deficient and Hexa/Neu3 doubly deficient mouse cells [39], total sialylation was flat (ω^2^ = 0.009, not significant) while the composition of α2-3 sialylation moved, as Neu5Gcα2-3Gal went from undetectable in controls to 7% and then 10% of the parent (redistribution p = 0.0032, Fig. 5E). The source study reported altered high-mannose and complex *N*-glycans [39] but made no claim about sialic-acid species, making this a new observation. It should not be read as a NEU3 mechanism, however. NEU3 is not restricted to α2-3 linkages [40] and prefers Neu5Ac over Neu5Gc [41], so a gain in Neu5Gc share is the opposite of what its substrate preference would predict, which is why we propose CMAH flux or lysosomal sialic-acid recycling as the more plausible explanations.

## Discussion

The single number we would ask a reader to carry away from our study is 45%. Nearly half of the motif-level findings that a standard analysis would report, and that a paper would list in its abstract, carry no independent signal, as they are the shadow of a change one or more levels down. This is not an argument against motif analysis, which remains the only tractable readout of a non-templated polymer. It is an argument that a motif result is incomplete without its residual, in the same way that a gene-set result is incomplete without knowing whether the child sets drive it.

The complementary number is 138, the magnitude of our significant redistribution findings. Because the balances are reference-free, redistribution is detectable exactly where marginal analysis is weakest, at abundant, near-universal motifs whose totals cannot move much. The *N*-glycan core nodes that dominate this regime are structurally obligate and therefore have almost no marginal dynamic range, which is precisely why their internal reorganization typically goes unreported.

The approach we developed here is also not specific to released glycans. Any measurement whose features are ordered by containment (e.g., site-level glycoproteomics, lectin-array motifs, glycan-binding specificities) admits the same graph, the same stratified correction, and the same residual, as we have shown with the example of site-level glycoproteomics.

Limits are worth stating plainly. Redistribution is a statement about co-occurrence structure, not about biosynthesis. A parent whose children shift has changed the contexts in which it appears, which constrains but does not identify the enzymatic cause, and GlyCompare’s precursor ordering answers the complementary question. The containment order is only as informative as the motif vocabulary that generated it, so a sparse or badly chosen feature set yields a shallow graph and little decomposition. The two-group balance test requires fewer balances than samples, which is satisfied for 91% of parents here but might represent a limitation in very small studies.

Placed in the wider field, this work sits at the meeting point of two lines that have not previously been connected. Comparative glycomics has been steadily professionalized with motif quantification and differential abundances [19], compositional treatment of relative abundances [20], AI-driven structure assignment from tandem mass spectra [42,43], regular-expression matching over glycan sequences [44], and biosynthetic network construction [45,46]. Yet in all of it the motif table has remained a flat list. Meanwhile the graph-structured multiple-testing literature [21–23] matured around gene sets and was never ported, because glycomics lacked an ordering that was both automatically derivable and quantitatively enforced. Containment supplies exactly that, and the abundance inequality it imposes is what upgrades the port from an analogy into an exact decomposition.

Three directions follow naturally. The first is a proper contest against hierarchical FDR, focus-level, and elim-style pruning, adapted to a data-derived graph. Our comparison is against the flat correction the field uses, which establishes the practical gain but might not yet represent the optimal graph-aware procedure that future work could work towards. The second is to connect redistribution to enzymology. A parent whose children reapportion is describing the outcome of competition between glycosyltransferases for a shared acceptor, which is precisely the regulatory layer that non-templated synthesis makes hard to observe [11]. Pairing redistribution events with transcript or activity measurements for the implicated transferases would convert a description into a mechanism, and the disialyl-T antigen convergence reported here, with its two redundant synthases, is the obvious first test. The third, as already alluded to above, is scope. Containment orders exist for glycoproteomic compositions, as we show, but equally for motifs in glycan arrays and lectin arrays [47], predicted binding specificities [48], and the substructures used by glycan machine-learning models [49], where the same redundancy quietly inflates feature importance.

Finally, the practical consequence for readers of the glycomics literature is concrete. When a study reports that sialylation or galactosylation has changed, that statement is, in our corpus, more likely than not an echo of a change at a more specific structure, and that specific structure is both more interpretable and more actionable. Reporting the residual alongside the marginal costs nothing, since both are already produced by the same analysis, and will aid in driving more concrete and actionable insights into modulations of the glycome in health and disease.

## Methods

### Datasets

All 45 comparative glycomics datasets in the glycowork (v1.10.0) glycomics data loader [50] were used for this work (30 two-group, 15 multi-group; median 14 samples, range 6 to 286), spanning *N*-glycans, *O*-glycans, and glycosphingolipids across human and mouse tissues, cell lines, and body fluids. No dataset was excluded for any reason.

### Motif quantification and graph construction

Motifs were quantified with the “exhaustive” (all occurring mono– and disaccharides, as well as their wildcarded subsumptions) and “known” (152 literature-known motifs of biological relevance) feature sets from glycowork. Motifs with fully identical abundances (e.g., Sia and Neu5Ac, if Neu5Ac was the only sialic acid in a dataset) were deduplicated so that the least ambiguous label survived, with string length used only to break ties among equally ambiguous labels. Containment edges were tested by termini-aware subgraph isomorphism using glycowork, prefiltered by abundance dominance, oriented by ambiguity with a positional-index fallback, and transitively reduced (all contained in the *glycowork*.*motif*.*annotate*.*get_motif_dag* function).

### Formal definitions

Let *M* be the set of motifs quantified in a dataset and *a*_*m*_(*s*) the abundance of motif *m* in sample *s*. Containment ⊑ is a partial order on *M*, and *m* ⊑ *m′* implies *a*_*m*_(*s*) ≥ *a*_*m*_′(*s*) for all *s*. The graph is the transitive reduction of ⊑, so that ch(*p*) = {*c*: *p* ⊑ *c* and no *m* with *p* ⊑ *m* ⊑ *c*}. For a parent *p* we define the residual

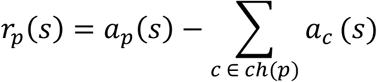

the part of *p* occurring outside every child context, and the sub-composition *x*_*p*_(*s*) = (*a*_{*c*1}_(*s*), …, *a*_{*ck*}_(*s*), *r*_*p*_(*s*)), whose parts sum to *a*_*p*_(*s*) by construction. Its additive logratio balances with respect to the first part are

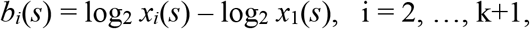

which are invariant to any per-sample scalar λ(*s*), since *b*_*i*_(λ*x*) = *b*_*i*_(*x*). This is why the balances require neither a reference component nor a scale model, in contrast to the marginal statistics, which are computed after an additive logratio transformation of the full table. Redistribution tests H_0_: E[b | group] is constant, by Hotelling’s T^2^ for two groups and by one-way PERMANOVA on Euclidean distances between balance vectors for more than two. The residual test is a univariate test on log_2_ *r*_*p*_(*s*) returned to the same logratio frame as the marginals.

### Sensitivity estimator

For a method *M* at effect-size window [*τ, κ*] let *R*(*τ*) be the number of motifs called significant with effect size at least *τ* and at most *κ*, where *κ* = 5 for Cohen’s *d* and is unbounded for ω^2^, which is already bounded by construction, and let *R*^*b*^(*τ*) be the same count on the b-th of *B* = 100 label permutations, under which no true effect exists. The estimated number of true positives then is

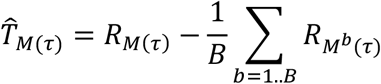

so that each method controls for its own false-positive rate before methods are compared. Per-dataset differences were tested with a two-sided sign-flip permutation test on the mean difference (50,000 sign assignments), stratified so that signs were permuted within the two-group and multi-group designs separately, since the two use different effect-size gates and are therefore on different scales. A rank-based test is inappropriate here because the improvement is strongly asymmetric in magnitude and gains are large and losses are near zero, a distinction signed ranks cannot represent. Robustness was assessed by leave-one-dataset-out and by a 10% winsorized variant of the same test.

### Testing

Two-group comparisons used moderated t-tests with Cohen’s *d*, and multi-group comparisons a moderated one-way ANOVA with ω^2^. Variance moderation follows the empirical-Bayes framework of Smyth [26]: each motif’s within-group variance is shrunk toward a prior, and because motifs that contain one another are measured on overlapping structures and therefore share measurement noise, that prior is the geometric mean of the variances over the motif’s own containment neighborhood (its parents and children in the DAG) rather than over all motifs. The prior degrees of freedom are estimated per dataset by Smyth’s moment estimator and capped at the residual degrees of freedom, so that the prior can contribute at most as much information as the data, in the spirit of the bounded estimation of Phipson et al. [27]. This cap matters, since an unconstrained estimate on a chance-homogeneous variance set produced p-values as small as 1e-32 from six samples. Reported effect sizes are computed from the unmoderated variance and are therefore unaffected. Where no containment graph exists, as in sequence-level analysis, the prior falls back to the global geometric mean. Motifs were assigned to the family of their rarest root ancestor and corrected by two-stage adaptive Benjamini–Hochberg within families. Redistribution used Hotelling’s T^2^ test on the sub-composition balances for two groups and a one-way PERMANOVA with 999 permutations for more than two. Residual, redistribution, and marginal p-values were each corrected as separate families.

### Benchmark

We compared our new approach with glycowork v1.9.0 that relies on flat multiple testing correction and an unmoderated test statistic. Both arms shared precomputed preprocessing, and each was run on 100 label permutations per dataset (sign-flips for paired designs). Default effect-size gates were a Cohen’s *d* of 0.5 and ω^2^ of 0.06 (except for the mentioned effect size gate sweep).

## Data availability

All data used in this article can either be found in supplemental tables or as stored datasets within glycowork.

## Code availability

Code and documentation are available via glycowork v1.10.0 (https://github.com/BojarLab/glycowork).

## Acknowledgement

This work was supported by the Swedish Foundation for Strategic Research and the University of Gothenburg, Sweden.

## Contributions

Conceptualization: D.B., Funding Acquisition: D.B., Resources: D.B. Software: D.B., X.Z., Supervision: D.B., Visualization: D.B., X.Z., Writing—Original Draft Preparation: D.B., X.Z., Writing—Review & Editing: D.B., X.Z.

