## Supplemental Figures for "Expressing glycan motifs as a containment order increases sensitivity and uncovers motif redistributions"

### Supplementary Figures

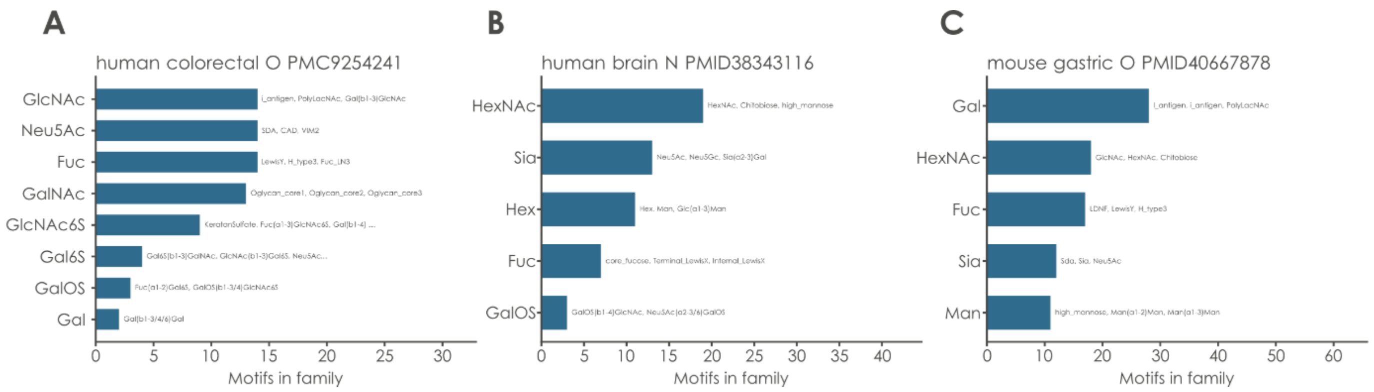


**Supplementary Figure 1. DAG families recapitulate biosynthetic branches.** For three representative datasets, every family containing more than one motif is shown as a bar whose length is the number of member motifs; the y-axis label is the family's root motif and three example members are printed inside each bar. Families are the grouping unit for the stratified multiple-testing correction.

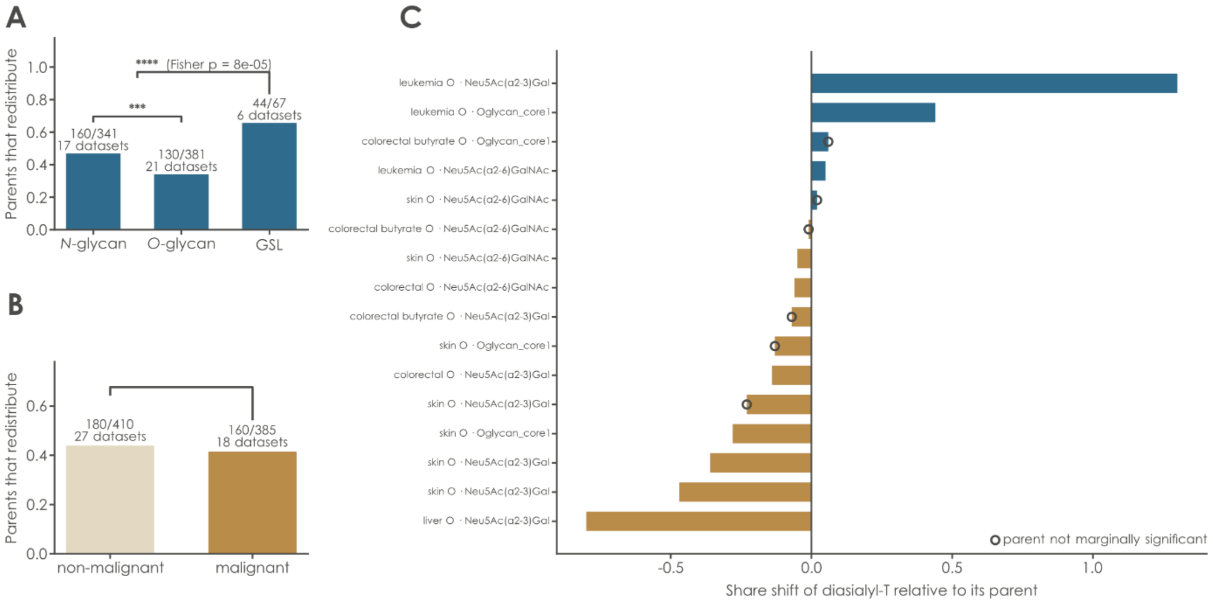


**Supplementary Figure 2. Analyzing where redistribution occurs. A)** Fraction of testable parent motifs that redistribute significantly, by glycan class; counts and dataset numbers are printed above each bar and brackets give two-sided Fisher exact tests. **B)** The same as (A), for malignant versus non-malignant datasets. **C)** Every significant redistribution event whose explaining child is the disialyl-T antigen, one bar per dataset and parent, signed by the direction of the disialyl-T share shift: blue, disialyl-T gains share; brown, it loses share. Open circles mark events where the parent motif is not itself marginally significant.

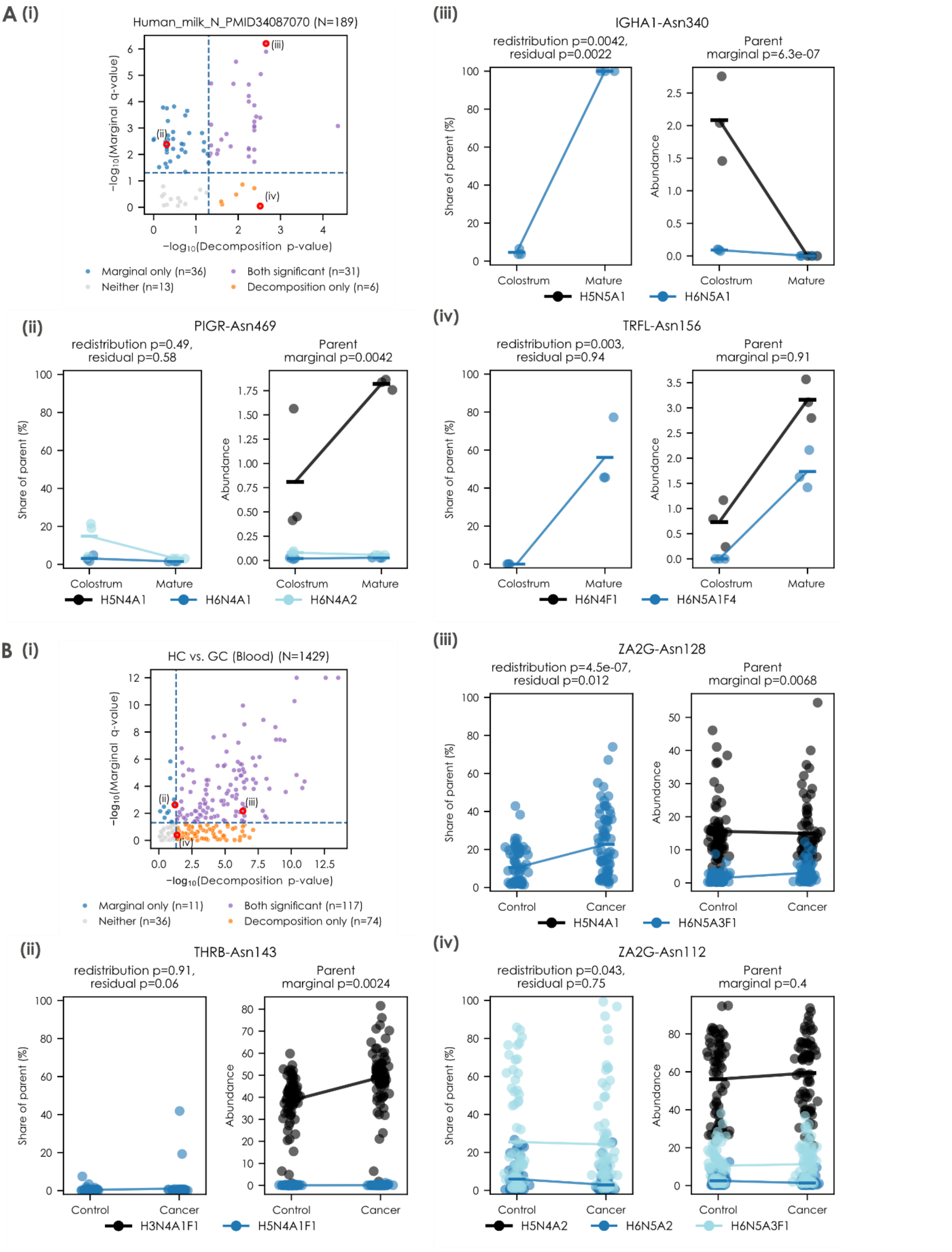


**Supplementary Figure 3. Site-specific glycan compositions exhibiting decomposition signals in colostrum versus mature milk (A) and healthy versus gastric cancer individuals' blood (B).** (i) Scatter plots showing the marginal q-value on the y-axis and the smaller of the residual and redistribution q-values on the x-axis; on a negative log10 scale, with the dash line at q = 0.05 (n: number of findings, N: number of component-wise composition containment). Site-specific compositions with significant (ii) marginal q-values only; (iii) both significant marginal q-values and decomposition q-values; (iv) decomposition q-values only.
